# Streamlining Multiplexed Tissue Image Analysis with PIPΣX: An Integrated Automated Pipeline for Image Processing and EXploration for Diverse Tissue Types

**DOI:** 10.1101/2025.05.04.652145

**Authors:** Mariya Mardamshina, Frederic Ballllosera Navarro, Anna Martinez Casals, Christophe Avenel, Carolina Wählby, Emma Lundberg

## Abstract

Spatial proteomics via multiplexed tissue imaging is transforming how we study biology, enabling researchers to investigate dozens of markers in a single tissue section and explore how cells behave in their native habitat. While imaging technologies have advanced rapidly, data analyses remain a bottleneck. To address this, we developed **PIPΣX** (Pipeline for Image Processing and EXploration), a user-friendly, end-to-end open-source software designed to make complex image analysis approachable, even for those with little or no programming skills. PIPΣX combines robust automation with an intuitive graphical user interface, guiding users through each step of the analysis, from image preprocessing and membrane-aware cell segmentation to signal quantification and spatial data exploration. Each feature includes built-in explanations, recommendations, and quality controls to help users make confident choices throughout the process. PIPΣX is compatible with a wide range of multiplexed imaging platforms, and its outputs integrate seamlessly with visualization tools like TissUUmaps and QuPath. Also, it supports downstream applications by enabling direct export of selected cell coordinates for laser microdissection. This functionality facilitates precise isolation of target cell populations for deep proteomic or transcriptomic profiling. With PIPΣX, researchers can extract meaningful biological insights from multiplexed images more easily and robustly, helping to bridge the gap between powerful imaging technologies and real-world scientific discovery.

**Highlights:**

- PIPΣX offers a user-friendly “one-stop shop” pipeline for multiplexed tissue image analysis
- without coding
- Works across diverse tissue types and imaging platforms at whole-slide scale
- Includes membrane-aware segmentation and quality control features
- Seamlessly integrates with visualization platforms like TissUUmaps and QuPath for data exploration
- Enables export for automated laser microdissection and spatial single-cell profiling

## Introduction

Recent advances in spatial proteomics have opened exciting new doors in biomedical research^1^. Using techniques like cyclic immunofluorescence, barcoded technologies (Phenocycler-Fusion/CODEX^2,3^), and metal conjugated antibodies (IMC^4^, MIBI^5^), scientists can now visualize dozens to hundreds of proteins in a single tissue section, at single-cell resolution, while preserving the spatial context that’s often crucial for understanding disease, development, and tissue organization^6,7^. This has led to major discoveries in fields ranging from cancer biology^8^ to neuroscience^9^. However, to analyze and interpret these rich, complex images is not easy. The analysis pipelines often rely on a combination of multiple software tools, manual parameter tuning, and programming experience, barriers that can be overwhelming for many researchers, especially those coming from experimental or clinical backgrounds.

While several tools exist for different parts of the analysis workflow, like CellProfiler^10^ for segmentation, QuPath^11^ for annotation, or Cellpose^12^ for deep-learning-based object detection, most of these require a fair amount of computational knowledge to set up and use effectively, or are limited to smaller image sizes. Some are optimized for specific tissue types or imaging platforms, while others offer flexibility, often at the cost of user-friendliness. Pipelines like MCMICRO^13^ and Squidpy^14^ provide powerful analysis environments but assume users are comfortable working in Python or using the command line. As a result, many scientists with access to advanced imaging technologies still struggle to fully analyze and interpret their data. To help bridge this gap, we developed **PIPΣX** - a fully integrated, easy-to-use “one-stop-shop” pipeline built specifically for researchers with extensive, minimal, or no prior computational experience. What sets PIPΣX apart is its focus on accessibility without sacrificing analytical power. From the beginning, we designed the tool to be navigable through a graphical user interface (GUI) that’s intuitive and informative. Each feature, whether it’s image preprocessing, segmentation, marker quantification, or clustering, comes with built-in explanations, example parameters, and interactive guidance. This makes it easier for users to understand what each step does and how to adjust settings to fit their own data.

PIPΣX also supports end-to-end workflows, from raw image input to interactive spatial visualization, and includes direct export to platforms like TissUUmaps for downstream exploration. It works with data from multiple multiplexed imaging modalities, offering a consistent and reproducible framework that doesn’t require switching between multiple tools or writing custom scripts.

In this work, we introduce the design and features of PIPΣX, and we demonstrate how it performs across a variety of use cases, highlighting its ability to make multiplexed tissue image analysis more transparent, interactive, and accessible to the broader scientific community.

## Results

PIPΣX is a Python-based software suite designed to convert raw or minimally processed immunofluorescence imaging data of any complexity (from low to high-plex projects) acquired directly from high-content imaging platforms into interpretable, analysis-ready datasets. The pipeline supports all major stages of image analysis: preprocessing, segmentation, quantification, clustering, visualization, and interactive data exploration. By integrating modular scripts into a unified interface, PIPΣX allows both novice and expert users to process large-scale imaging datasets with minimal effort and without prior knowledge of any programming language.

### PIPΣX Enables Fully Automated, End-to-End Multiplexed Image Analysis Across Diverse Tissue Types

Designed with accessibility in mind, PIPΣX allows users to process high-resolution, whole-slide images, from raw data to interpretable results, without requiring computational expertise or scripting. Its architecture supports integration with existing imaging platforms (e.g., Akoya, CODEX, MIBI) and formats (e.g., TIFF, QPTIFF, PNG, JPEG) as well as downstream analytical tools, providing a reproducible framework adaptable across tissue types and experimental settings (Fig. 1).

**Fig. 1:**
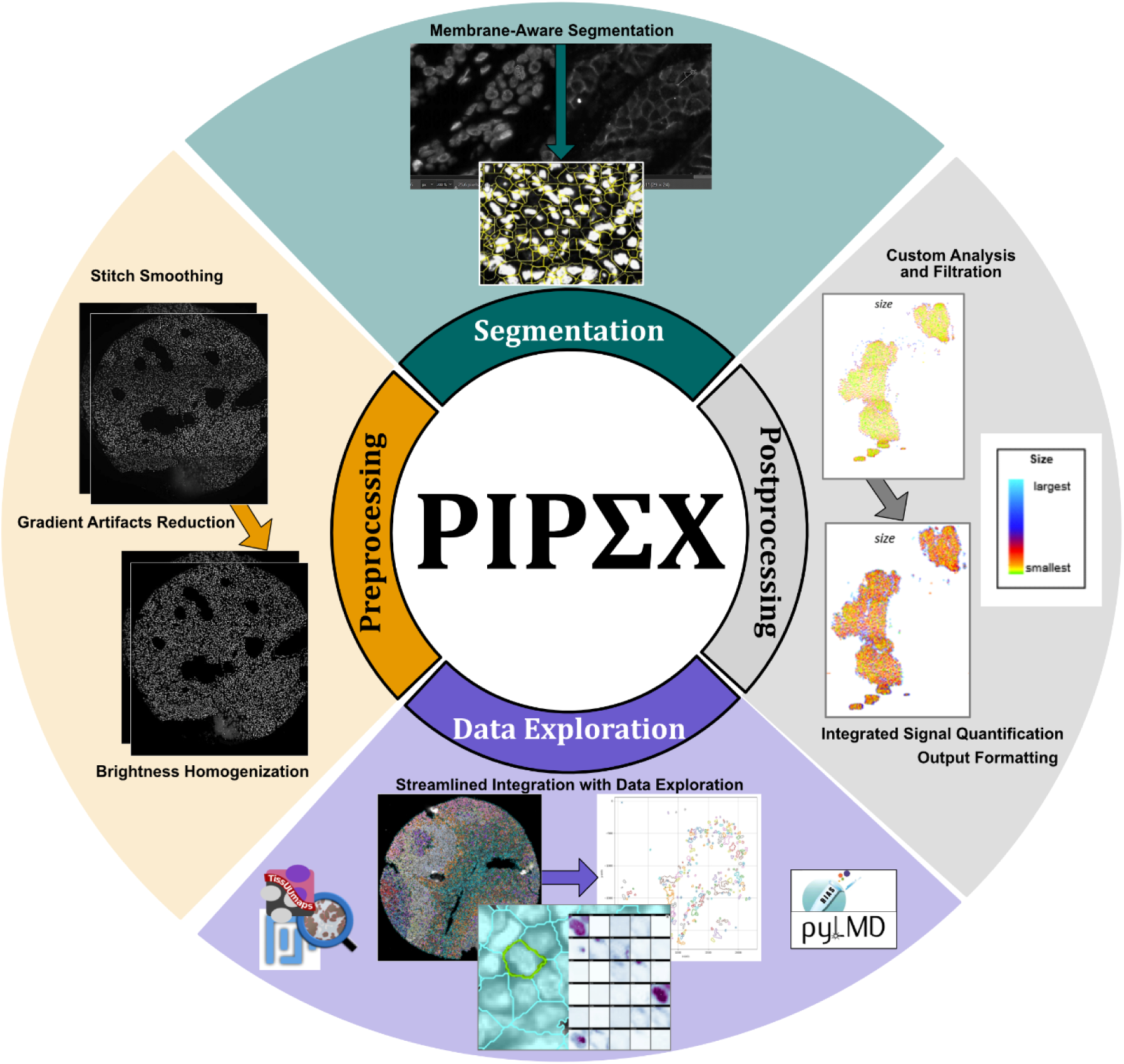
PIPΣX software capabilities and flexibility with the possibility of independent use of different modules. Overview of the PIPΣX pipeline, illustrating its modular design for flexible and scalable image analysis across diverse tissue types. Each component, preprocessing, segmentation, clustering, visualization, and export, can be run independently or as part of an end-to-end automated workflow. The modular structure supports both standalone use of individual steps and integration into broader spatial analysis pipelines, enabling customization for user-specific applications such as clustering, cell-type annotation, spatial context exploration, or export for laser microdissection (LMD).

To benchmark PIPΣX, we tested its performance across multiple datasets, including multiplexed images from pancreas, kidney, mammary gland, tonsil, and esophageal tissues. These images varied in staining complexity (ranging from 2 to 40+ channels), resolution, and sample preparation protocols. Across all datasets, PIPΣX consistently performed robust image preprocessing, accurate segmentation, and cell-wise quantification with minimal user intervention. The pipeline includes built-in quality control checkpoints and outputs structured data formats compatible with common analysis environments (e.g., scanpy, napari, and TissUUmaps)^15–18^.

### Robust Image Preprocessing Across Variable Inputs

PIPΣX introduces a modular preprocessing framework that corrects for a range of acquisition-related artifacts frequently encountered in multiplexed tissue imaging. Unlike existing tools that rely on hard-coded parameters or manual curation, PIPΣX leverages adaptive algorithms that operate efficiently on gigabyte-scale, whole-slide images, with configurable options for iterative refinement based on downstream visual or quantitative outputs. The preprocessing pipeline includes intensity thresholding, tile correction procedures, each of which can be applied independently or in combination (Fig. 2a–c).

**Fig. 2:**
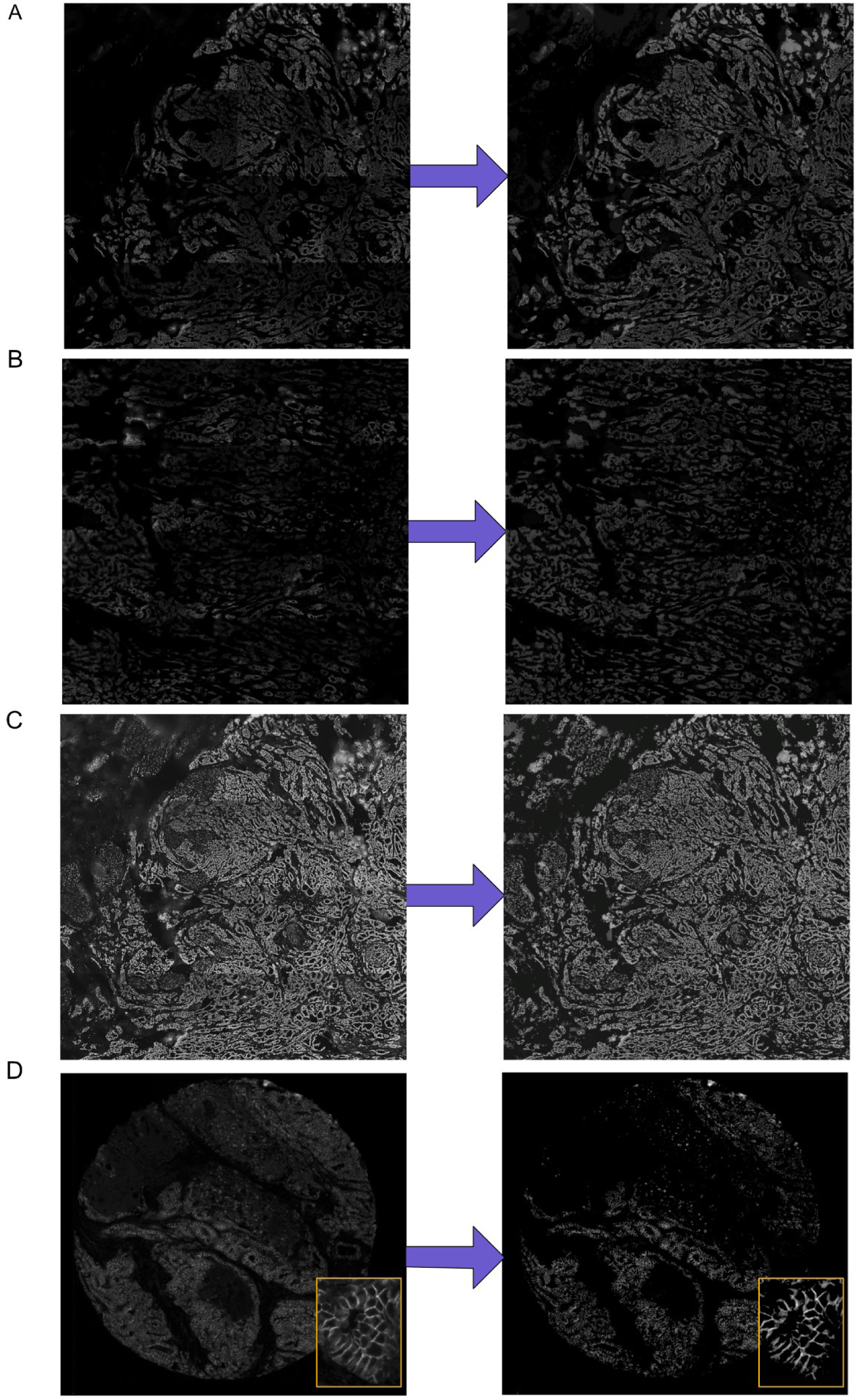
Preprocessing strategies enhance tissue image clarity for analysis in PIPΣX. Representative examples of raw and preprocessed multiplexed images demonstrate the impact of automated correction on image quality. A) Breast cancer whole-tissue section exhibiting uneven illumination and visible tile boundaries. Preprocessing parameters: threshold minimum = 1, bright levels = 5:1:2, light gradient correction = 2, exposure = 200. B) High background signal and pronounced tiling artifacts in breast cancer tissue.

Intensity thresholding enables the suppression of background noise and technical debris, such as folding artifacts or precipitates, by applying configurable lower and upper percentile cutoffs. This step allows users to retain only the most informative intensity ranges for analysis.

To further refine signal quality, PIPΣX implements multilevel Otsu thresholding, which automatically partitions the image into multiple intensity classes based on global histogram features. This approach enhances detection of biologically meaningful structures by identifying optimal intensity boundaries without user bias and is especially useful when intensity distributions are complex or multimodal. Users may define both the number of threshold levels and the subset of classes used for binarization or segmentation.

Tile correction addresses four major challenges introduced by tile-based image acquisition: gradient light reduction, brightness homogenization, and stitch smoothing. Gradient light reduction segments each tile into local kernels to detect and suppress directional illumination artifacts or vignetting. Brightness homogenization adjusts histogram distributions across tiles to reduce inter-tile variability and promote global intensity consistency. Stitch-smoothing minimizes abrupt transitions at tile boundaries, yielding seamless, whole-slide views. We benchmarked the preprocessing module across multiple imaging formats, including tissue microarray (TMA) cores and large whole-tissue sections. In all cases, preprocessing eliminated uneven lighting, harmonized brightness across tiles, and recovered structurally informative regions that were previously obscured by background contamination (Fig. 2d).

By addressing these acquisition-specific artifacts at the earliest stage of analysis, PIPΣX enables more robust segmentation and quantification, especially when working with archival or low-quality data.

### Flexible Segmentation with User-Defined or Built-In Options

PIPΣX includes a modular segmentation framework designed to accommodate a range of tissue architectures, experimental designs, and user expertise. The default pipeline begins with StarDist^19–21^, a deep-learning-based nuclei detection algorithm that performs well across many multiplexed imaging datasets. While effective for convex nuclear shapes, StarDist does not capture cytoplasmic or whole-cell morphology and can encounter memory constraints when applied to large-scale, high-resolution images.

To address these practical limitations, PIPΣX includes adjustments to reduce memory usage, such as using more efficient internal routines from StarDist’s source code and an optional Linux-specific swap memory mechanism. When memory limits are still exceeded, the pipeline automatically applies downscaling for segmentation, followed by upscaling for reintegration, maintaining alignment with subsequent analysis steps. We typically work with the standard format size of the Phenocycler-Fusion whole slide imaging platform, and the area is 3.5 cm x 1.5cm.

Beyond nuclear detection, PIPΣX supports an optional expansion step to approximate full-cell boundaries. This method relies on user-defined parameters to control object growth and minimize overlap, providing a simple means of estimating cytoplasmic extent when no membrane marker is available.

When membrane-localized channels are included in the dataset, the pipeline optionally applies membrane-aware watershed segmentation to further refine cell boundaries. This approach incorporates basic morphological priors (e.g., compactness, size constraints) and may improve boundary accuracy in epithelial or densely packed tissues.

The final segmentation output can combine nuclear and membrane-based masks. In this step, membrane regions are associated with corresponding nuclei and intersected with expanded nuclear objects to generate more complete cell representations. While not intended to provide perfect morphological reconstructions, this approach can improve cell delineation in certain contexts, particularly where nuclear overlap or irregular spacing challenges conventional methods.

We compared different segmentation strategies using representative normal and cancerous tissues from diverse organ sites (Fig. 3). To improve cell boundary delineation, PIPΣX optionally incorporates membrane signal into segmentation through a membrane-aware refinement step. The choice of pan-membrane marker is guided by the biological context of the study: for epithelial-rich solid tumors, broadly expressed epithelial junctional markers, e.g., E-cadherin (CDH1), Beta Catenin1 (CTNNB1) or Pan-cytokeratin cocktail (PanCK) are typically selected; in contrast, for immunologically focused tissues such as tonsil or bone marrow, hematopoietic markers like CD44 or CD45 offer better membrane definition for immune cell segmentation, as demonstrated in our comparative tissue analysis.

**Fig. 3:**
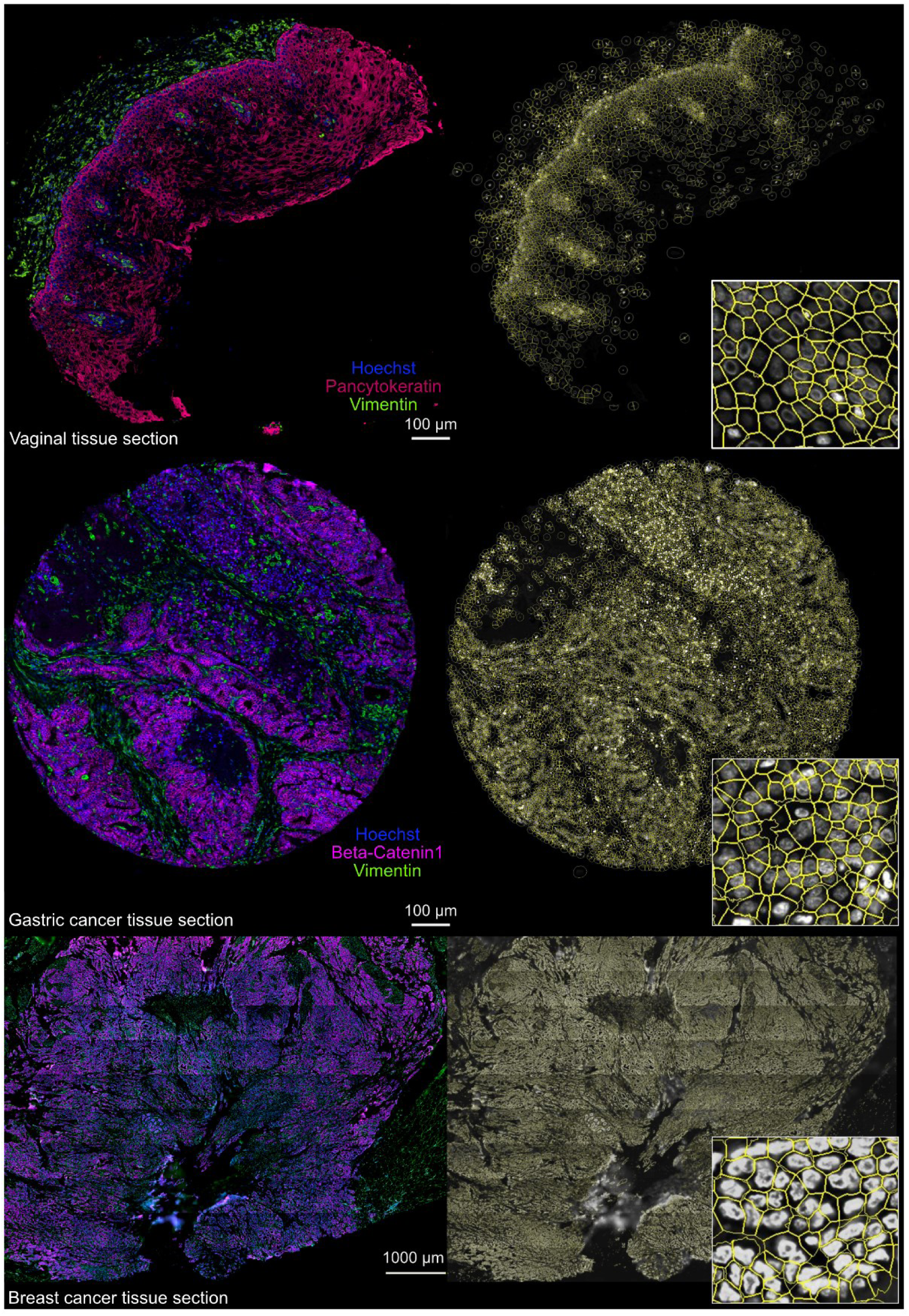
PIPΣX segmentation with membrane refinement captures cellular detail across tissue types and states. Examples from multiple tissue types demonstrate accurate cell segmentation using nuclear and membrane cues, with optional refinement for enhanced cellular boundary resolution. **Top panel**, Vaginal tissue section segmented using StarDist on Hoechst-stained nuclei. Pan-cytokeratin staining was used for membrane-aware refinement. The panel includes a raw image, a segmentation mask, and a zoom-in illustrating the precise delineation of individual cells. **Middle panel**, Gastric cancer tissue section segmented with StarDist. Membrane refinement was performed using β-Catenin1 staining, improving boundary fidelity in densely packed epithelial regions. **Bottom panel**, Close-up view of a representative field showing Hoechst and β-Catenin1 staining along with the corresponding segmentation mask in breast cancer tissue section, highlighting the improvement in membrane contour definition following refinement. C) Another example is from breast cancer tissue with unbalanced tiles and a diffuse background. Preprocessing with adjusted light gradient correction (level = 3) improves uniformity. D) Gastric cancer core showing high background autofluorescence. Preprocessing with bright levels = 5:2:4 and exposure = 100 enhances signal-to-noise and clarity.

Additionally, depending on tissue type or state, PIPΣX includes customizable settings for nuclear segmentation. This allows users to define the proximity of cells to each other and adjust the definition of nuclei, which is especially useful for irregularly shaped or overlapping nuclei. These settings enhance segmentation accuracy in challenging tissue morphologies, providing better resolution for boundary delineation in complex tissues.

StarDist alone yields clean nuclear masks but does not reflect cytoplasmic or membrane marker patterns (Fig. 4a). Expansion-based segmentation, commonly used in tools such as QuPath or Halo AI, provides broader cell approximations but can introduce artifacts due to overlapping cells.

**Fig. 4:**
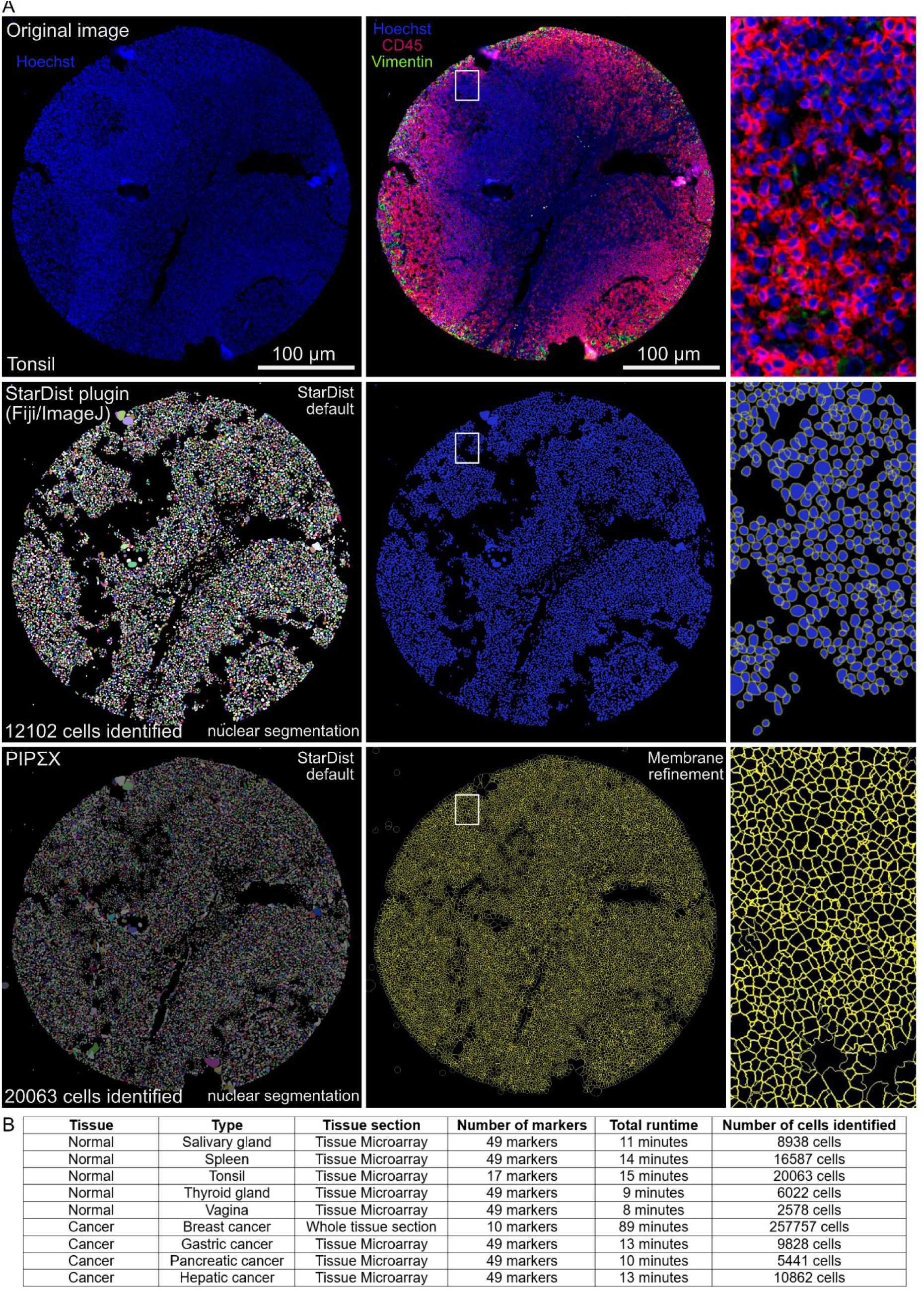
Impact of membrane refinement on segmentation performance in tissue images. Comparison of segmentation outcomes with and without membrane-aware refinement in a normal, non-diseased tonsil section. A) **Top panel:** Original images used for segmentation. Hoechst staining identifies nuclei in both Fiji and PIPΣX workflows, while CD45 membrane staining is used for refinement in PIPΣX. **Middle panel:** Segmentation results using StarDist with default settings (Fiji/ImageJ), followed by watershed-based expansion. A zoomed-in view highlights limitations in accurately resolving cell boundaries. **Bottom panel:** PIPΣX segmentation using StarDist paired with membrane refinement based on CD45 staining. The resulting segmentation mask demonstrates improved cell border definition, particularly in densely clustered immune cells. A close-up view is shown on the right. B) Summary table of the PIPΣX performance across diverse tissue types and biological states. The table reports the number of cells segmented and total runtime per image, highlighting the scalability and consistency of the PIPΣX workflow. Tissue types include both normal and diseased states, spanning epithelial and immune-rich environments.

The combined approach in PIPΣX reduces some of these limitations by incorporating membrane signal information where available, though performance varies depending on tissue morphology and staining quality.

To support segmentation assessment, the pipeline includes visual quality control modules that overlay results on the original image alongside summary statistics such as the membrane refinement confidence score. The score can be calculated by interpreting the number of non-zero values in the membrane refinement column in the csv output table, which indicates if the cell shape has been affected by the membrane detection. These tools allow users to evaluate segmentation outcomes prior to quantification, enabling informed adjustments to parameters if needed.

Recognizing the diversity of available tools and workflows, PIPΣX also allows users to import external segmentation masks. These can be generated using algorithms such as Cellpose^12^, DeepCell^22^, SAM^23,24^, or custom-trained models and incorporated at various stages of the pipeline. This flexibility enables researchers to tailor segmentation strategies to their specific use case without needing to bypass or re-implement other parts of the analysis pipeline.

The segmentation process ends with the single-cell image marker signal quantification. For every cell, the pipeline computes four key metrics: the mean marker intensity, the 90th percentile of non-zero pixel intensities (capturing local expression hotspots), the fraction of cell pixels above the threshold (indicating spatial enrichment), and 3 bins histogram thresholding coefficients (via Otsu thresholding) that quantifies the separation between marker signal and background. These features are collated into a structured data table that retains spatial coordinates, enabling downstream phenotyping, comparative analysis across tissue types, and integration with tools such as TissUUmaps or Python-based workflows.

This approach is critical for distinguishing subtle yet biologically relevant expression differences in complex tissues. For example, it enabled the detection of polyhormonal endocrine populations in pancreatic islets and the delineation of progenitor-enriched niches in renal tissue^25^. Notably, metrics such as the 90th percentile intensity are particularly effective for identifying low-abundance, spatially restricted signals, such as granule-localized hormones or transcription factors, that may be missed by conventional mean-based quantification.

Users can choose to modify the pipeline at different stages, for example, extracting nuclear features only or running full-cell segmentations, depending on their requirements. This modularity is designed to accommodate diverse analytical goals while maintaining compatibility with downstream quantification and spatial analysis.

The whole workflow is quite robust, and a typical run-through takes approximately 10 minutes from start to finish for a tissue core around 10.000 cells and a bit more than an hour for a whole tissue section with around 260000 cells (Fig. 4b).

### Integrated Signal Quantification, Filtration, and Cluster Refinement

As an initial step for the downstream analysis, PIPΣX provides several options to seamlessly include simple and common mathematical data processing steps like normalization and batch correction, all with a simple parameter toggle.

To support quality control and downstream biological interpretation, PIPΣX also includes an optional marker-based filtration module. This feature is designed to remove potential artifacts such as over-segmented nuclei or edge effects, which can bias population-level statistics. As part of the pipeline’s configurable settings, users can choose to exclude cells falling within the top 1% of intensity for nuclear markers (DAPI or Hoechst) and epithelial markers (CDH1 and CTNNB1), helping to filter out potential artifacts such as under-segmentation, tissue folds, or edge effects. To ensure accurate operation, marker identifiers in the dataset must match these default names precisely.

To facilitate interactive data review and hypothesis generation, the pipeline offers tools to explore marker coexpression patterns (Fig. 5a). This includes pairwise coexpression and other plots as well as gating tools to assess overlapping marker distributions across spatially distinct cell subsets, aiding in the identification of mixed phenotypes or transitional states. In parallel, PIPΣX supports interactive visualization of marker expression surfaces in three dimensions, enabling users to explore expression heterogeneity in x, y, and z space by overlaying multiple markers within the same spatial context (Fig. 5b). This capability highlights regional variation and rare expression patterns that might be missed in two-dimensional plots, improving contextual interpretation of spatially organized biology.

**Fig. 5:**
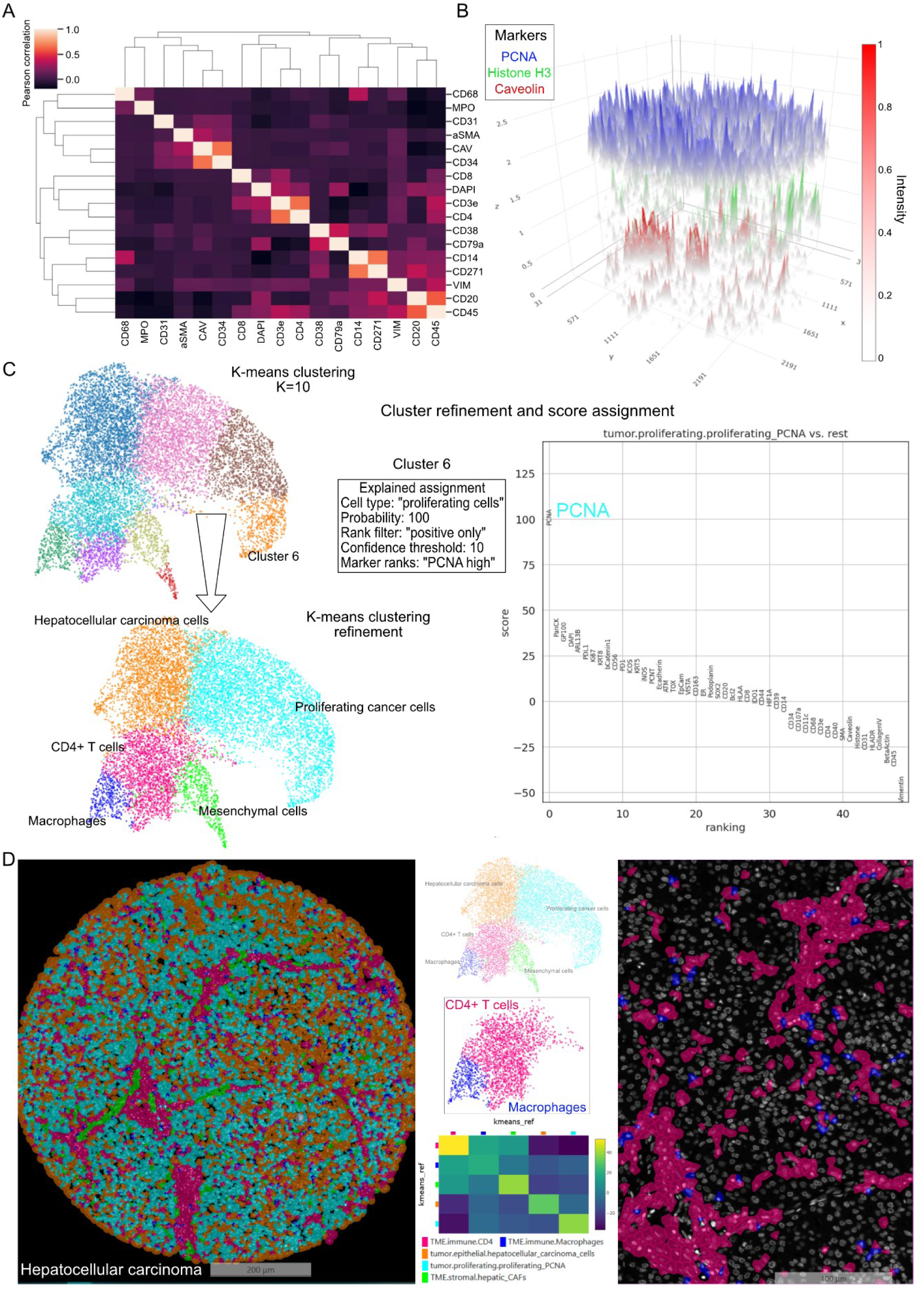
Comprehensive tissue analysis with PIPΣX: plotting, annotation, and neighborhood exploration. PIPΣX enables integrative analysis of spatial expression data, supporting dimensionality reduction, clustering, and spatial context mapping. A) Heatmap of marker co-expression across cells in a non-diseased tonsil sample. Pearson correlation highlights relationships among immune and structural markers. B) Surface marker plots illustrating expression levels of PCNA, Histone H3, and Caveolin in spatial coordinates (x, y, z), visualizing spatially resolved molecular gradients of marker expressions. C) **Top:** UMAP embedding of all segmented cells color-coded by K-means clustering output (K = 10). **Bottom:** Refined clustering output generated using PIPΣX cluster refinement algorithm, with assignment scores shown in **the right panel** based on marker-specific contributions. D) **Left:** Segmentation mask in tissue space, color-coded by refined cluster identities. **Middle:** UMAP and corresponding neighborhood heatmap highlighting spatial proximity between macrophages (blue) and CD4⁺ T cells (magenta) based on Euclidean distances. **Right:** Spatial localization of these two clusters in tissue mirrors the proximity relationships observed in the heatmap.

For datasets requiring deeper phenotypic resolution, PIPΣX also incorporates a cluster refinement procedure that improves the interpretability of unsupervised clustering results (e.g., Leiden, K-means). This module enables users to define biologically meaningful cluster annotations using a flexible, rule-based system (Fig. 5c; top panel). Cluster labels are refined by comparing ranked marker expression profiles within each cluster to user-defined expectations. The algorithm assigns each cluster the most appropriate annotation based on a confidence score that quantifies how closely the expression profile matches the specified rules (Fig. 5c; bottom panel). Users can further customize the behavior through parameters such as rank_filter (e.g., using only positively expressed markers) and min_confidence (threshold for rule adherence), allowing for stringent or permissive refinement depending on experimental goals.

Cluster labels are assembled from three user-specified components (cell_group, cell_type, and cell_subtype), enabling consistent and hierarchical annotation across datasets. This design supports both manual review and high-throughput curation of complex spatial datasets, where unsupervised clusters may not always map clearly to known biological cell types.

Together, these features establish PIPΣX as a robust and extensible framework for image-based single-cell analysis. Its multi-metric quantification, modular architecture, and user-guided refinement tools make it well-suited for diverse applications in spatial biology, from exploratory tissue profiling to targeted hypothesis testing.

#### Seamless Integration with External Image Viewers Enables Interactive Data Exploration Visualizing Segmentation Results with QuPath

PIPΣX easily integrates with QuPath, a popular software used in digital pathology, to help users visualize and explore their segmentation results. When you run PIPΣX with the generate_geojson command, a cell_segmentation_geo.json file is created in your analysis folder. This file can be imported into QuPath to view the segmented cells. The process is simple: start a new project in QuPath and add your images. Then, open the script editor in QuPath and copy the example script provided with PIPΣX. Update the script to point to the cell_segmentation_geo.json file. This is followed by running the script to visualize your data in QuPath.

This integration works with QuPath version 0.3. Future QuPath updates may require adjustments to the import process.

#### Interactive Visualization and Quality Control of Spatial Proteomics Data via TissUUmaps Integration

To facilitate interactive data exploration and visual quality control of high-plex spatial proteomics datasets, PIPΣX provides native integration with TissUUmaps, an open-source web-based viewer optimized for large-scale tissue imaging data. By enabling the generate_tissuumaps option, users can export an .h5ad file containing spatial coordinates, segmentation metadata, marker intensities, and clustering results. This file can be directly loaded into TissUUmaps for immediate visualization, and an optional web-export (include_html=yes) generates a self-contained version that can be hosted or shared as an interactive resource in any browser.

One of the most critical features supported by this integration is the dynamic overlay of cell segmentation masks on any reference channel, such as DAPI or specific protein markers. This functionality allows users to assess segmentation accuracy rapidly and identify imaging or staining artifacts, such as misaligned markers, signal bleed-through, or cell boundary errors, before performing downstream analyses. To support interpretability and cluster refinement, the segmentation masks can be color-coded by any categorical label, including unsupervised clustering results (e.g., K-means, Leiden) or curated cell type annotations.

To further enhance exploratory analysis, we integrated and adapted a suite of TissUUmaps plugins originally developed for spatial transcriptomics, tailoring them for spatial proteomics applications in the PIPΣX pipeline (Fig. 6a):

**Fig. 6:**
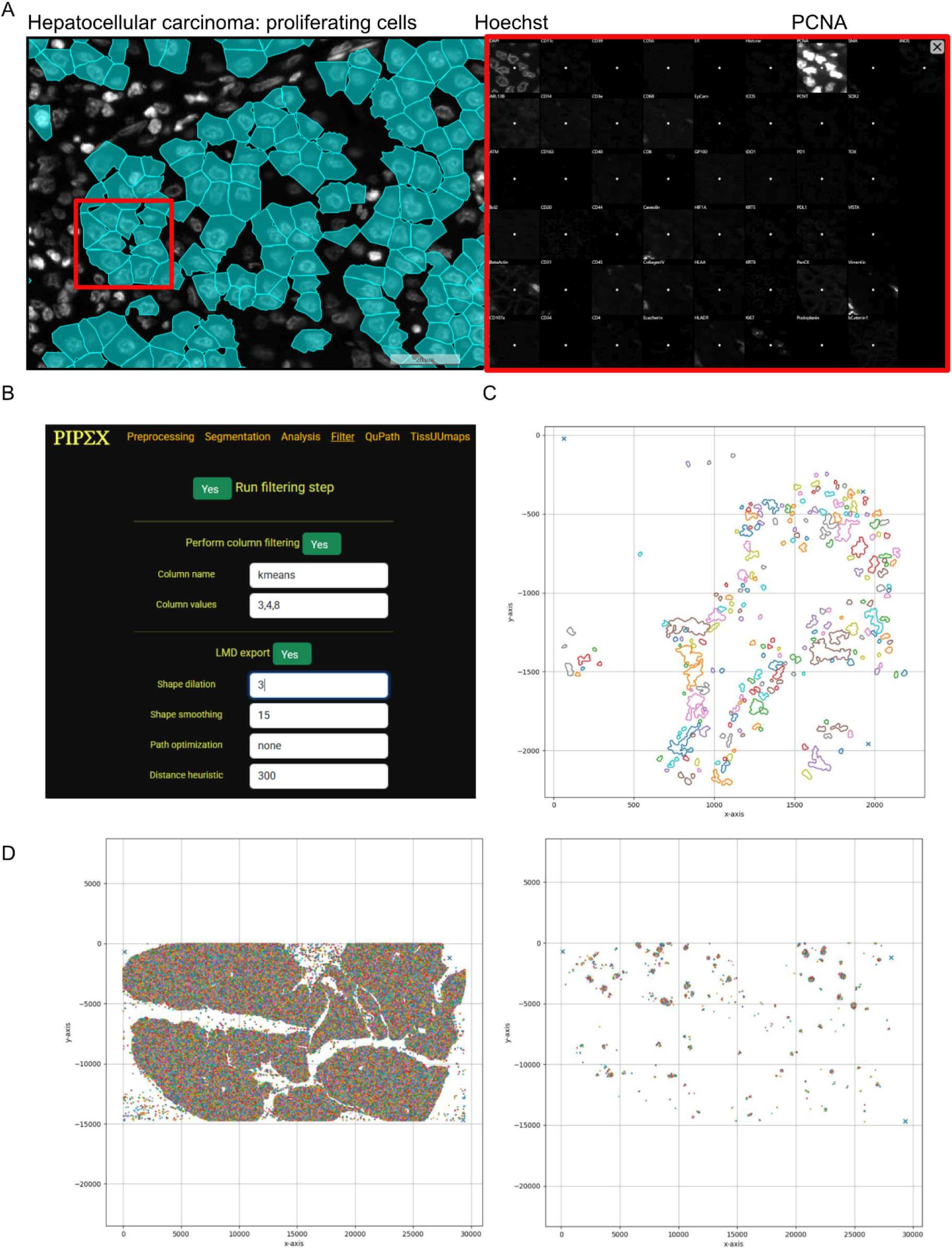
Interactive image review and export of defined cell clusters for LMD applications. PIPΣX supports integration with interactive visualization tools and downstream workflows such as laser microdissection (LMD). A) Visualization of the segmentation mask of hepatocellular carcinoma tissue using the TissUUmaps Spot Inspector plugin. Cluster of proliferating cells selected from PIPΣX output is interactively overlaid onto the original image. **Right:** Close-up views of selected areas highlight marker expression patterns consistent with cluster annotations. B) Schematic of the pipeline enabling automated export of cell contours for LMD. PIPΣX provides directly usable annotation outputs compatible with microdissection systems. C) Scatterplot showing contours selected for export from a non-diseased tonsil tissue section. Highlighted cells correspond to a user-defined cluster of interest. D) **Left:** Full scatterplot displaying all segmented cells included for export. **Right:** Subset of selected cells for targeted microdissection. Calibration points used for LMD alignment are marked with crosses, corresponding to automated placements generated by PIPΣX.

The Spot Inspector plugin, initially designed as a quality control tool for image-based spatial transcriptomics, was modified to support spatial proteomics data. It allows users to click on any point in the tissue image and instantly open a gridded subview displaying signal intensities across all imaging rounds and channels at that location. Barcoded trace overlays highlight corresponding molecular signatures, enabling efficient validation of combinatorial labeling schemes and detection of decoding or segmentation errors. This is particularly useful for distinguishing ambiguous or transitional cell states.

The Feature Space plugin was adapted to visualize low-dimensional embeddings (e.g., UMAP, PCA) generated during PIPΣX analysis. Originally developed to explore transcriptomic profiles, it now supports proteomic data without requiring additional processing. Users can interactively select cell groups within the feature space and immediately highlight their spatial distribution on the tissue image. This functionality supports rapid interrogation of phenotypic heterogeneity and spatial organization of rare or functionally distinct cell populations.

The InteractionV&QC plugin, designed for neighborhood-based analyses in spatial transcriptomics, was similarly extended to work with proteomic data. It enables visualization of spatial cell–cell interactions derived from PIPΣX’s integrated neighborhood analysis module. Validated cell classifications can be used to generate interaction heatmaps, reflecting enrichment or depletion of proximity-based relationships across clusters or annotations. These tools allow users to inspect whether observed spatial patterns reflect underlying biology or technical variation.

Finally, for figure generation and image export, the Scale Bar plugin provides precise and customizable scale bar overlays. This ensures visual consistency across datasets and facilitates the creation of high-quality, annotated figures suitable for publication.

Together, these capabilities make the PIPΣX-TissUUmaps integration a powerful, interactive environment for spatial proteomics, enabling users to validate data quality, explore cellular heterogeneity, and communicate results with clarity and precision.

### Integration of pyLMD for Single-Cell Contour Export and Multiomics Profiling

To enable precise downstream analysis of rare or biologically relevant cell populations, we integrated the open-source pyLMD library into the PIPΣX pipeline. This integration facilitates the seamless export of single-cell contours for laser microdissection (LMD), ensuring accurate targeting of user-selected cells based on high-dimensional image analysis. The ability to extract contours of interest directly from segmented images is essential for spatially resolved multiomics workflows, such as deep proteomics or transcriptomics, especially when profiling rare cell states.

Key functionalities from pyLMD have been adapted and extended to allow fine control over contour generation, including automatic assignment of calibration coordinates required for LMD software compatibility. This automation eliminates the need for manual calibration steps and minimizes user error during laser capture setup. Importantly, our implementation supports multiplexed calibration within a single tissue section, enabling simultaneous acquisition of distinct regions or phenotypes for parallel multiomics readouts.

To support reproducibility and transparency, the pyLMD export process is fully automated and traceable. As illustrated in Fig. 6b-d, the export workflow proceeds through a series of computational steps: (1) flagging LMD export during segmentation, (2) identifying calibration points, (3) defining the image transformation matrix, (4) applying shape dilation to ensure laser precision, (5) optimizing cutting paths, and (6) merging segmented regions prior to XML export. This process ensures compatibility with Leica LMD systems and allows researchers to iteratively refine cutting regions based on evolving annotations.

By enabling precise and flexible export of contours from annotated tissue images, this feature maximizes tissue utility and minimizes sample waste, allowing multi-modal profiling from a single section. This makes PIPΣX especially well-suited for studies with limited sample availability or those requiring integration of spatial context with high-resolution molecular data.

### Additional features are available to adopt an image format for analysis

To facilitate broader adoption and compatibility of PIPΣX with diverse imaging platforms, we have included auxiliary scripts for image pre-processing in the Cell Profiling GitHub repository (https://github.com/CellProfiling/ell_code_template/tree/master/examples). These utilities are designed to streamline the conversion and preparation of multiplexed imaging datasets prior to PIPΣX execution. Specifically, the scripts support the extraction of individual tissue regions and the conversion of complex formats, such as .qptiff, into single .tif files suitable for downstream analysis. These resources enable users to preprocess images originating from high-content imaging systems like the Akoya Vectra Polaris and CODEX, thereby extending PIPΣX’s interoperability with established experimental pipelines.

## Discussion

The ability to visualize dozens to hundreds of proteins in tissue sections has opened new opportunities in understanding how cells behave in health and disease. Yet for many researchers, especially those without programming experience, analyzing these complex images remains a major challenge. With PIPΣX, we set out to make advanced multiplexed image analysis more approachable while still providing the tools needed to generate meaningful scientific insights.

A key strength of PIPΣX is its intuitive graphical interface. Rather than asking users to write code or navigate complicated file structures, the software guides them step-by-step through image cleaning, cell segmentation, signal measurement, and interactive data exploration. Built-in tips, quality checks, and recommended settings help users understand each step and make confident decisions. This makes PIPΣX especially valuable for biologists, pathologists, and clinicians who want to explore their data directly without relying on specialized bioinformatics support.

Another important feature is the controlled environment in which the software runs. Because PIPΣX manages all tools and settings behind the scenes, users can be confident that their analysis is consistent and reproducible, even across different labs or studies. This is particularly useful for large teams or clinical settings, where standardization is essential.

In addition to making analysis easier, PIPΣX helps users go deeper. It supports the interactive exploration of spatial relationships between cell types using tools like TissUUmaps and QuPath and enables downstream applications such as laser microdissection for single-cell proteomics or transcriptomics. These features allow researchers to move seamlessly from imaging to molecular profiling, opening new possibilities for discovery in tissues with rare or complex cell populations. Making PIPΣX available as a web-based tool presented an opportunity to make it easier to use in resource-limited settings or as part of shared computing environments in research centers or even hospitals.

While PIPΣX offers many strengths, there are still areas for improvement. For example, cell segmentation quality depends on the clarity of certain stains, such as those that highlight cell membranes. In the future, we plan to add features that let users correct segmentation errors more easily and improve the software’s ability to learn from such feedback or combine several markers to improve the membrane completeness and make the staining fully pan-membrane. Looking ahead, we envision expanding PIPΣX to support new types of data, such as spatial transcriptomics and histopathology, and to include tools for automated pattern recognition, phenotype prediction, and cell type prediction based on the neighborhood associations. These features would help users uncover subtle features or rare cell types that may be difficult to identify manually.

In summary, PIPΣX lowers the barriers to analyzing multiplexed tissue images by offering a streamlined, user-friendly interface that combines automation with flexibility. It empowers researchers from a range of backgrounds to engage with their data directly, promoting more transparent, reproducible, and insightful spatial biology research. As imaging technologies continue to evolve, tools like PIPΣX will play an essential role in making spatial analysis accessible to the broader scientific community.

## Methods

### Overview of Pipeline Structure

PIPΣX (Pipeline for Image Processing and Exploration) is a modular, GUI-driven software framework developed in Python for the automated analysis of multiplexed tissue images (https://github.com/CellProfiling/pipex). It supports the complete image analysis workflow from raw image input to downstream data exploration, including preprocessing, segmentation, quantification, clustering, and visualization. The pipeline operates within a controlled software environment and is deployable across local and institutional computing systems. Image preparation steps, including the conversion of .qptiff files to individual .tif images and the extraction of single tissue crops from TMAs, were performed using utility scripts available at https://github.com/CellProfiling/ell_code_template/tree/master/examples.

### Image Preprocessing

To address acquisition-related artifacts commonly present in whole-slide, multiplexed immunofluorescence images, PIPΣX includes a dedicated preprocessing module. This module provides a suite of configurable operations designed to improve image quality by correcting background noise, light gradients, and tiling inconsistencies. All preprocessing steps can be applied independently or in combination, and intermediate results are saved in JPEG format for visual inspection.

### Intensity Thresholding

Minimum and maximum intensity thresholds (-threshold_min, -threshold_max) are used to remove background fluorescence and high-intensity artifacts such as tissue folds or precipitates. These thresholds are defined as percentiles of the global intensity distribution and allow for rapid suppression of outliers while preserving signal within the informative range.

### Otsu-Based Binning and Filtering

Users can optionally define multi-level Otsu thresholding (-otsu_threshold_levels) to classify intensity values into discrete segments. Thresholding can be applied either with a default setting (e.g., 3, corresponding to three intensity classes with default bin filtering of 1:2) or with explicit bin specifications (e.g., 5:1:2), which can be used to generate masks or remove specific intensity bands from the original image.

### Tile Correction and Homogenization

To mitigate illumination inconsistencies and visible tile boundaries in large mosaic images, the module includes several corrective steps:

**Gradient Light Correction** (-light_gradient): Each tile is subdivided into smaller kernel patches, and local normalization is applied to correct for directional illumination artifacts and vignetting. Gradient complexity can be tuned with a value from 1 to 4, where higher values allow for finer-grained correction.

**Brightness Homogenization** (-balance_tiles): Tile-to-tile intensity variation is reduced through histogram-based scaling and standard deviation comparisons, harmonizing the brightness distribution across the entire image.

**Stitch Smoothing** (-stitch_size): The interface regions between adjacent tiles are smoothed over a user-defined pixel width to eliminate visible seams and improve continuity.

### Exposure Rescaling

The final preprocessed image can be globally brightened or dimmed using the -exposure parameter, which rescales pixel intensities relative to the original image. Thi is offered as a convenient (and optional) quality of life final step to avoid the need for an intensity rescaling in other visualizing software.

### Example Configurations

The module has been used to successfully process diverse sample types with a range of acquisition challenges: Light gradients and uneven tile intensities were corrected using - threshold_min=1, -otsu_threshold_levels=5:1:2, -light_gradient=2, and -exposure=200.

In cases with strong background and pronounced tiling artifacts, the same parameters were sufficient to suppress noise and enhance contrast.

For more challenging gradients and poorly balanced tiles, a stronger correction (- light_gradient=3) and histogram normalization (-balance_tiles=yes) were applied.

Autofluorescent tissues were handled using more selective intensity binning (- otsu_threshold_levels=5:2:4) and moderate exposure boosting (-exposure=100).

Together, these preprocessing options enable users to flexibly salvage and improve low-quality images, supporting accurate segmentation and robust downstream analysis.

### Cell Segmentation

Cell segmentation in PIPΣX is initiated using StarDist, a deep-learning algorithm for nuclei detection based on DAPI/Hoechst signals. The default pipeline includes nuclear segmentation using StarDist. Optional cell expansion from nuclei masks to approximate whole-cell morphology and membrane-aware refinement, activated when membrane marker channels are provided, using a watershed-based approach constrained by morphological priors. To manage large, whole-slide images, the pipeline includes automatic downscaling and reintegration steps for memory efficiency. Final segmentation outputs are saved as labeled NumPy arrays, and JPEG previews of each stage are generated for visual quality control.

External segmentation masks (e.g., from Cellpose^26^, DeepCell^22^, or CellSAM^24^) can be imported into the pipeline at user-defined stages. This ensures compatibility with custom models and varied segmentation strategies.

### Signal Quantification and Feature Extraction

Quantitative feature extraction is performed for each segmented cell using the following steps: marker image normalization and multi-level Otsu thresholding, per-cell metric computation, including mean marker intensity, 90th percentile intensity of non-zero pixels, fraction of pixels above threshold, and normalized Otsu separation score. These features are compiled into a structured CSV table with spatial coordinates. Optional marker-based filtration excludes outlier cells (e.g., top 1% of DAPI/CDH1/CTNNB1 intensities) to reduce segmentation artifacts.

### Unsupervised Clustering and Annotation Refinement

The pipeline supports unsupervised clustering (e.g., K-means, Leiden) and integrates a rule-based refinement module for biological annotation. Clusters are matched to user-defined marker expression rules and assigned hierarchical labels (cell group, type, subtype) based on a confidence score. Parameters such as rank_filter and min_confidence are user-adjustable to control annotation stringency.

### Export and Interactive Visualization

PIPΣX supports the automatic generation of multiple output formats: FCS files with per-cell data for flow-style visualization or external analysis tools, and GeoJSON overlays for QuPath visualization. The .h5ad files for TissUUmaps integration, including segmentation masks, marker intensities, cluster annotations, and spatial coordinates. HTML exports for browser-based interactive exploration using custom plugins (e.g., Spot Inspector, Feature Space, InteractionV&QC). All results are organized into a standardized output directory structure for reproducibility and ease of sharing.

### Batch Processing

For multi-sample workflows, PIPΣX supports batch-mode processing. Each sample can be configured with its own preprocessing, segmentation, and analysis settings, enabling parallel processing of large, heterogeneous datasets (e.g., multiple CODEX panels or tissues, TMAs).

### Laser Microdissection Export (pyLMD Integration)

PIPΣX includes an embedded module based on the open-source pyLMD library for laser microdissection (LMD) targeting. This feature allows users to export selected cell contours as LMD-compatible XML files for downstream proteomics or transcriptomics. The export process includes the identification of calibration points, shape dilation for precise contour cutting, transformation and optimization of laser paths, and XML file generation with merged and labeled targets. This integration supports multiplexed calibration and iterative refinement of cut regions within a single section.

## Code availability

All code used for image analysis and pipeline execution in this study is publicly available on GitHub. The PIPΣX software, including documentation and example workflows, can be accessed at https://github.com/CellProfiling/pipex. Additional utility scripts to facilitate image preprocessing, such as conversion of .qptiff files to single .tif and extraction of individual tissue crops, are available in the Cell Profiling code template repository at https://github.com/CellProfiling/ell_code_template/tree/master/examples. These tools support the integration of PIPΣX with common imaging formats and enhance accessibility for users working with diverse imaging platforms.

## Acknowledgements

We gratefully acknowledge funding from the EU Horizon 2020 programme (ESPACE 874710), Erling-Persson (HDCA), Knut and Alice Wallenberg Foundation (HDCA), HIRN (NIH 1U01DK120447), and The Chan Zuckerberg Initiative. This work was delivered as part of the PROMINENT team supported by the Cancer Grand Challenges partnership funded by Cancer Research UK (CGCATF-2021/100010), the National Cancer Institute (OT2CA278713), and the Scientific Foundation of the Spanish Association Against Cancer, AECC. We thank Frida Björklund and Anna Bäckström for performing experiments and generating image data included in this study. We are also grateful to Dr. Dita Gratzinger at the Department of Pathology at Stanford University and Dr. Cecilia Lindskog Bergström at Uppsala University for providing tissue samples. We are grateful to all the members of the Lundberg group for fruitful discussions. We also acknowledge support from the BioImage Informatics Facility (NBIS Sweden), with funding from SciLifeLab and the National Microscopy Infrastructure NMI (VR-RFI 2019-00217).

## Competing interests

E.L. is an advisor for the Chan-Zuckerberg Initiative Foundation, Element Biosciences, Cartography Biosciences, Pfizer, and Pixelgen Technologies.

## Author contributions

AMC and FBN conceived the project and designed the software goals. FBN developed the software. AMC, MM, and EL deployed the software in Emma Lundberg’s lab. MM and AMC expanded PIPΣX’s analysis components and utilities. CA and MM integrated TissUUmaps into PIPΣX. CA and CW deployed the software in Carolina Wählby’s lab. MM wrote the paper, all authors edited and commented on the manuscript. EL supervised the project.

## References

1. Method of the Year 2024: spatial proteomics. Nat. Methods 21, 2195–2196 (2024).

2. Schürch, C. M. et al. Coordinated Cellular Neighborhoods Orchestrate Antitumoral Immunity at the Colorectal Cancer Invasive Front. Cell 182, 1341–1359.e19 (2020).

3. Black, S. et al. CODEX multiplexed tissue imaging with DNA-conjugated antibodies. Nat. Protoc. 16, 3802–3835 (2021).

4. Giesen, C. et al. Highly multiplexed imaging of tumor tissues with subcellular resolution by mass cytometry. Nat. Methods 11, 417–422 (2014).

5. Angelo, M. et al. Multiplexed ion beam imaging of human breast tumors. Nat. Med. 20, 436– 442 (2014).

6. Quail, D. F. & Walsh, L. A. Revolutionizing cancer research with spatial proteomics and visual intelligence. Nat. Methods 21, 2216–2219 (2024).

7. Xu, Y. et al. Multimodal single cell-resolved spatial proteomics reveal pancreatic tumor heterogeneity. Nat. Commun. 15, 10100 (2024).

8. Radtke, A. J. et al. Multi-omic profiling of follicular lymphoma reveals changes in tissue architecture and enhanced stromal remodeling in high-risk patients. Cancer Cell 42, 444–463.e10 (2024).

9. Piyadasa, H. et al. Multi-omic landscape of human gliomas from diagnosis to treatment and recurrence. *bioRxiv* 2025.03.12.642624 (2025) doi:10.1101/2025.03.12.642624.

10. Carpenter, A. E. et al. CellProfiler: image analysis software for identifying and quantifying cell phenotypes. Genome Biol. 7, R100 (2006).

11. Bankhead, P. et al. QuPath: Open source software for digital pathology image analysis. Sci. Rep. 7, 16878 (2017).

12. Stringer, C., Wang, T., Michaelos, M. & Pachitariu, M. Cellpose: a generalist algorithm for cellular segmentation. Nat. Methods 18, 100–106 (2021).

13. Schapiro, D. et al. MCMICRO: a scalable, modular image-processing pipeline for multiplexed tissue imaging. Nat. Methods 19, 311–315 (2022).

14. Palla, G. et al. Squidpy: a scalable framework for spatial omics analysis. Nat. Methods 19, 171–178 (2022).

15. Wolf, F. A., Angerer, P. & Theis, F. J. SCANPY: large-scale single-cell gene expression data analysis. Genome Biol. 19, 15 (2018).

16. Chiu, C.-L., Clack, N., & the napari community. napari: a Python Multi-Dimensional Image Viewer Platform for the Research Community. Microsc. Microanal. 28, 1576–1577 (2022).

17. Solorzano, L., Partel, G. & Wählby, C. TissUUmaps: interactive visualization of large-scale spatial gene expression and tissue morphology data. Bioinformatics 36, 4363–4365 (2020).

18. Pielawski, N. et al. TissUUmaps 3: Improvements in interactive visualization, exploration, and quality assessment of large-scale spatial omics data. Heliyon 9, e15306 (2023).

19. Schmidt, U., Weigert, M., Broaddus, C. & Myers, G. Cell Detection with Star-Convex Polygons. in Medical Image Computing and Computer Assisted Intervention – MICCAI 2018 (eds. Frangi, A. F., Schnabel, J. A., Davatzikos, C., Alberola-López, C. & Fichtinger, G.) 265–273 (Springer International Publishing, Cham, 2018). doi:10.1007/978-3-030-00934-2_30.

20. Weigert, M., Schmidt, U., Haase, R., Sugawara, K. & Myers, G. Star-convex Polyhedra for 3D Object Detection and Segmentation in Microscopy. in 2020 *IEEE Winter Conference on Applications of Computer Vision (WACV)* 3655–3662 (IEEE, Snowmass Village, CO, USA, 2020). doi:10.1109/WACV45572.2020.9093435.

21. Weigert, M. & Schmidt, U. Nuclei instance segmentation and classification in histopathology images with StarDist. in 2022 *IEEE International Symposium on Biomedical Imaging Challenges (ISBIC)* 1–4 (2022). doi:10.1109/ISBIC56247.2022.9854534.

22. Bannon, D. et al. DeepCell Kiosk: Scaling deep learning-enabled cellular image analysis with Kubernetes. 505032 Preprint at 10.1101/505032 (2020).

23. Kirillov, A. et al. Segment Anything. Preprint at 10.48550/arXiv.2304.02643 (2023).

24. Israel, U. et al. CellSAM: A Foundation Model for Cell Segmentation. 2023.11.17.567630 Preprint at 10.1101/2023.11.17.567630 (2025).

25. Mardamshina, M. et al. Multiplexed Deep Visual Proteomics Unveils Spatial Heterogeneity and Rare Endocrine States in Human Adult Pancreatic Islets. 2025.04.27.650857 Preprint at 10.1101/2025.04.27.650857 (2025).

26. Stringer, C. & Pachitariu, M. Cellpose3: one-click image restoration for improved cellular segmentation. Nat. Methods 22, 592–599 (2025).

